# Stimulation modulates cell assemblies linked with gene networks in the human temporal cortex *ex vivo*

**DOI:** 10.1101/2025.06.25.661589

**Authors:** H. Moore, M. Dehnad, A. Freelin, B. Granger, S. Subramanian, A. Kulkarni, S. Berto, B. C. Lega, G. Konopka

## Abstract

Deep brain stimulation of the temporal cortex can enhance learning and memory in the face of cognitive impairment. Despite the potential of such therapies, the neural and genetic mechanisms underlying the effect of stimulation on human brain circuits are not understood. To explicate direct mechanisms of neural modulation elicited by brain stimulation, we developed an *ex vivo* approach utilizing microelectrode array stimulation and recording of resected temporal cortex from neurosurgical patients. We find that stimulation preferentially increases firing rates in pyramidal cells compared to interneurons and also strengthens cell assemblies. Using single cell multiomics, we link the observed physiological changes to cell type-specific gene expression patterns. We detail gene regulatory networks that indicate preferential involvement of specific excitatory neuron subtypes and the response of non-neurons. We conclude that the overall impact of stimulation on the human temporal cortex is activation of specific excitatory neurons and enhanced cell assembly activity, and that these changes are supported by gene networks involving immediate early, synaptic, and ion channel genes. Our findings establish a foundation to identify targetable cell type-specific genetic signatures that may be harnessed for therapeutic benefit in future neuromodulation strategies.

## Introduction

Electrical stimulation of subcortical brain structures is a mainstay of therapies for movement disorders like Parkinson’s disease. Deep brain stimulation (DBS) has more recently been applied to neuropsychiatric conditions, namely treatment-resistant obsessive compulsive disorder and depression [1, 2]. Novel neuromodulation strategies targeting the temporal lobe, including the lateral temporal cortex [3–5], seek to restore memory function using brain stimulation [6, 7]. However, the absence of a mechanistic understanding of how exogenous stimulation alters brain circuit activity impedes the ability to develop new therapies for cognitive disorders, improve existing approaches, and design non-invasive treatments modeled on the effects of brain stimulation.

An emerging model of how stimulation impacts neuronal function is that stimulation modifies brain circuits through mechanisms of synaptic and circuit plasticity [2, 8]. Insights into the underlying molecular mechanisms of brain stimulation are derived from studies in rodents. For example, in mice, stimulating the fornix for 45 minutes resulted in upregulation of immediate early genes (IEGs) and genes related to synaptic plasticity and neuronal survival [9]. This builds on data from rat primary cortical neuron culture that delineated cell type-specific induction of activity-regulated genes (ARGs), such as *ARC, EGR1,* and *FOS,* after neuronal activation [10, 11]. Many ARGs encode transcription factors that support important neuronal functions including synaptic plasticity and neuronal assembly dynamics [12, 13]. Thus, current studies in non-human systems support a framework by which stimulation leads to gene expression changes that underlie the ability of neurons to form, maintain, and rewire synapses and circuits.

Neural circuits, especially in the temporal lobe, engage assemblies, organized groups of neurons that cooperatively fire on short time scales, to implement plastic representations and neural sequences [14, 15]. The formation of neuronal assemblies is a key physiological mechanism in brain circuits that underlie diverse cognitive functions and behaviors [14]. Theoretical and empirical evidence suggest that the gamma timescale (∼25 ms) is the optimal window in which to analyze assemblies [14, 16, 17]. Direct brain recordings in humans have identified such gamma-linked neuronal assemblies in the temporal lobe during episodic memory behavior [17]. In that study, successful memory formation was predicted by assembly flexibility by way of neuron ‘drift’, or the recruitment of new assembly member neurons, and through the strengthening of member co-activation specifically during memory encoding epochs. Assembly activity and flexibility following a stimulus can provide a model for local circuit plasticity that can also be linked with gene expression signatures. For example, evidence from rodents shows that specific neuronal populations are primed to participate in learning and memory [18, 19], and that the predisposition to join assemblies during memory formation is determined by transcriptional and epigenetic profiles that are influenced by cell activation states [13]. However, the dynamic genetic mechanisms that support neural circuit components in the human brain remain essentially unknown, especially as they pertain to brain stimulation. Although some studies have described the cellular changes that occur in tissue surrounding DBS electrodes in human post-mortem tissue [20, 21], it is impossible to use cadaveric studies to understand the effect of neuromodulation on assembly activity and concomitant differential gene expression because of the timescale in which these changes occur.

We therefore sought to address these outstanding ambiguities linked with neuromodulation therapies using a novel experimental approach in the human temporal cortex *ex vivo*. We hypothesized that stimulating temporal cortex slices with a microelectrode array would enhance cell type-specific gene expression networks that support local cell circuits through assembly activation and flexibility. We therefore profiled these genomic dynamics using an unbiased, genome-wide multiomic approach at the level of individual cells and measured neuronal assemblies using high resolution multi-electrode arrays. Our data support models by which stimulation alters local cell circuit activity. Moreover, the identification of gene expression signature changes in these cell types upon stimulation represent molecular targets for future pharmacological therapies that support the salubrious circuit modulation achievable through invasive stimulation.

## Results

### Stimulation preferentially increases activity in cells with pyramidal-like characteristics

We hypothesized that stimulation would alter local neuronal circuits predicated on cell type-specific gene expression changes. We first tested whether stimulation altered neuronal activity at the level of individual cells. We completed 37 stimulation experiments with brain tissue slices from 12 subjects (**Fig 1a; Supp. Table 1**). Tissue viability was demonstrated by the generation of spontaneous neural activity. An analysis of basic activity metrics across all electrodes in each recording showed a significant increase in the mean 90^th^ percentile firing rate after stimulation (Wilcoxon, p = 0.011, n = 29; **Fig 1b**). There was no difference in average spike amplitude (t-test, t(28) = 0.3723, p = 0.71; **Supp Fig 1b**) or percent active electrodes (Wilcoxon, p = 0.86, n = 29; **Supp Fig 1b**). Thus, our *ex vivo* preparation and stimulation conditions induced robust and consistent activity across samples.

**Figure 1.**
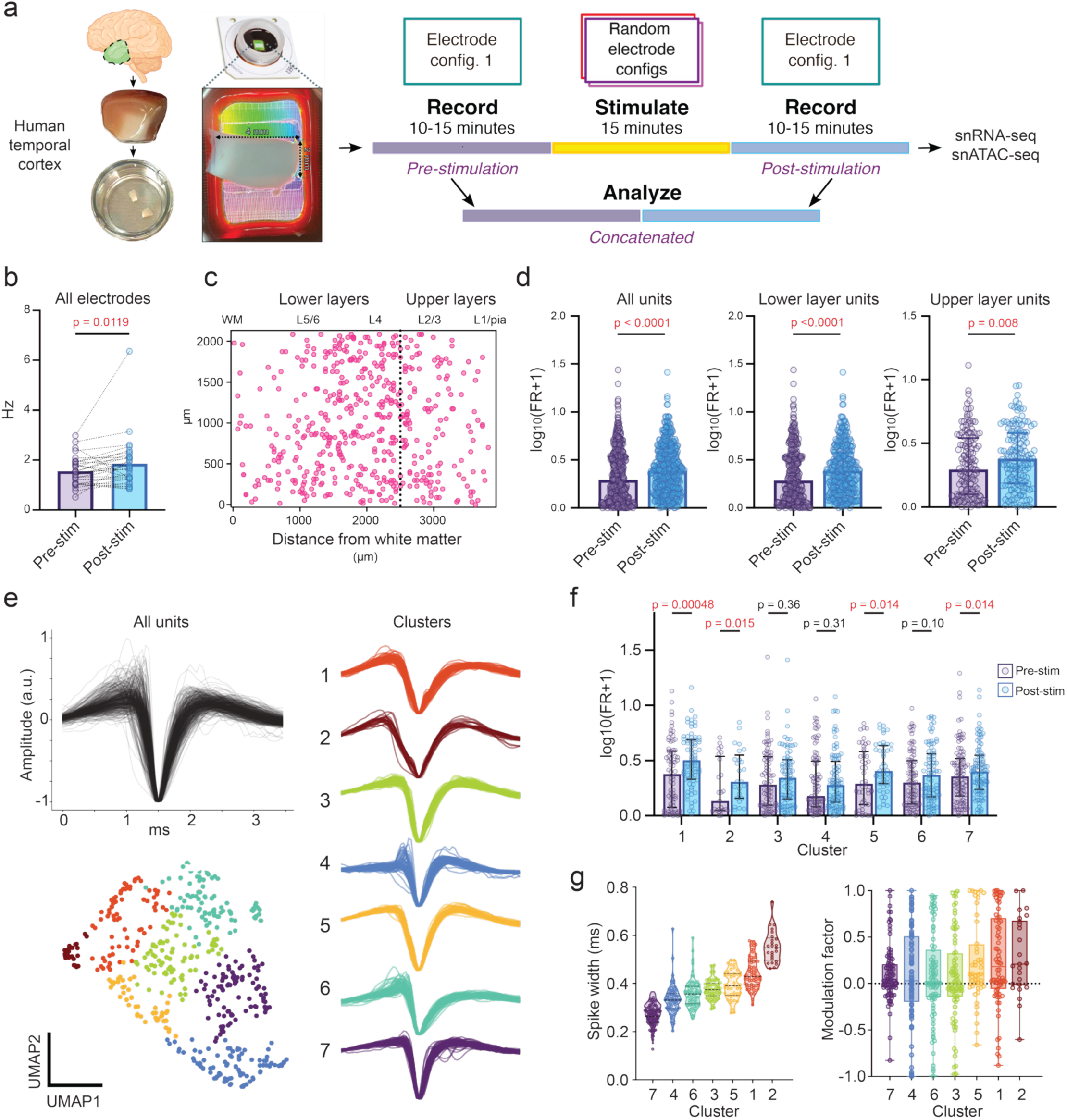
MEA stimulation increases firing rate activity in a cell-type dependent manner. **a.** MEA stimulation experiment overview. Left panel: a preparation of cortical slices from human temporal lobectomy samples. Right panel: stimulation experiment timeline. **b.** The mean 90^th^ percentile firing rate recorded at all electrodes increases after stimulation (Wilcoxon signed-rank test p=0.011*, n* = 29 experiments) **c.** The location of the highest amplitude spike of every unit identified with spike sorting. The boundary between the upper and lower layers is set at 1500 uM from the right short edge of the MEA, which corresponds to the pial edge of the slice. **d**. Stimulation increases the average firing rate of all units together (paired two-tailed Wilcoxon signed-rank test *p*<0.0001*, n* = 494 units), and in upper- (paired two-tailed Wilcoxon signed-rank test *p =* 0.008, *n* = 142 units) and lower-layer groups (paired two-tailed Wilcoxon signed-rank test *p*<0.0001*, n* = 352 units). **e.** Putative cell types identified with WaveMAP clustering. Upper left panel shows all unit waveforms that were averaged, aligned, trimmed, and normalized prior to clustering into groups in the right panel. **f.** Effect of stimulation on firing rate in each cell-type cluster. All comparisons were made with paired two-tailed Wilcoxon signed-rank tests. **g**. Unique characteristics of putative cell type clusters. Violin plot shows the distribution of spike trough widths at half maximum for each WaveMAP cluster. Bar and strip plot shows the firing rate modulation factor according to the calculation MF = (FRpost-FRpre)/(FRpost+FRpre). Clusters are ordered according to increasing spike width to show the positive relationship between this arrangement and the change in firing rate. In **d** and **g,** log10(FR+1) transform of firing rate values is for visualization only. Statistics performed on non-transformed data.

We next aimed to understand how stimulation affects individual neuron types. We identified 494 putative neurons and classified them according to approximate cortical layer location (**Fig 1c; Supp.** Fig 1c-d). The mean firing rate of all cells significantly increased after stimulation (Wilcoxon, p<0.0001, n = 494), an effect that was also apparent in upper- (Wilcoxon, p = 0.008, n = 142) and lower-layer cells (Wilcoxon, p<0.0001, n = 352) separately (**Fig 1d**). There was no difference in the average change in firing rate between upper- and lower-layer cells across the duration of the recording epoch (Mann-Whitney U, p = 0.43).

We then clustered the units into seven putative cell types based on the shape of each unit’s average spike waveform to test our hypothesis that the response to stimulation would differ between cell types (**Fig 1e**). Cells within each cluster were uniquely distributed across the cortical layers and had group-defining characteristics like trough width, duration, and peak to trough ratio (**Fig 1e-g; Supp.** Fig 1f-g). The average firing rate increased after stimulation in four of the seven cell clusters (Wilcoxon, FDR corrected p values = 0.00048, 0.015, 0.36, 0.31, 0.014, 0.10, 0.014 for Clusters 1 to 7, respectively; **Fig 1f**). Clusters with narrow spikes (mean spike width <0.35 ms – clusters 4 and 7), representing putative interneurons, responded less robustly to stimulation than clusters of putative pyramidal cells with broad spikes (mean spike width >0.35 ms – clusters 1, 2, 3, and 5). We found a positive correlation between firing rate modulation factor and trough width that could not be explained by chance (Spearman rank correlation, r = 0.104, p = 0.021; permutation test, p = 0.007; **Fig 1g**). These results suggest that units with characteristics similar to pyramidal cells are more sensitive to stimulation than units similar to interneurons.

### Neural assemblies are active in the human temporal cortex *ex vivo*

We employed a well-validated method for assembly identification that uses a combination of principal and independent component analyses to extract co-firing relationships at the 25 ms scale, with iteration of the ICA step to ensure rigor of identified assemblies (**Fig 2a**; **Supp.** Fig 2a) [22]. We identified 19 assemblies each consisting of independent, functionally connected member neurons (**Fig 2b**; **Supp.** Fig 2b). This exceeded the number of assemblies expected by chance in every iteration of the shuffle control procedure we implemented (permutation test, p<0.0001; see methods; **Supp.** Fig 2c). These results demonstrate that assemblies were present in our tissue samples.

**Figure 2.**
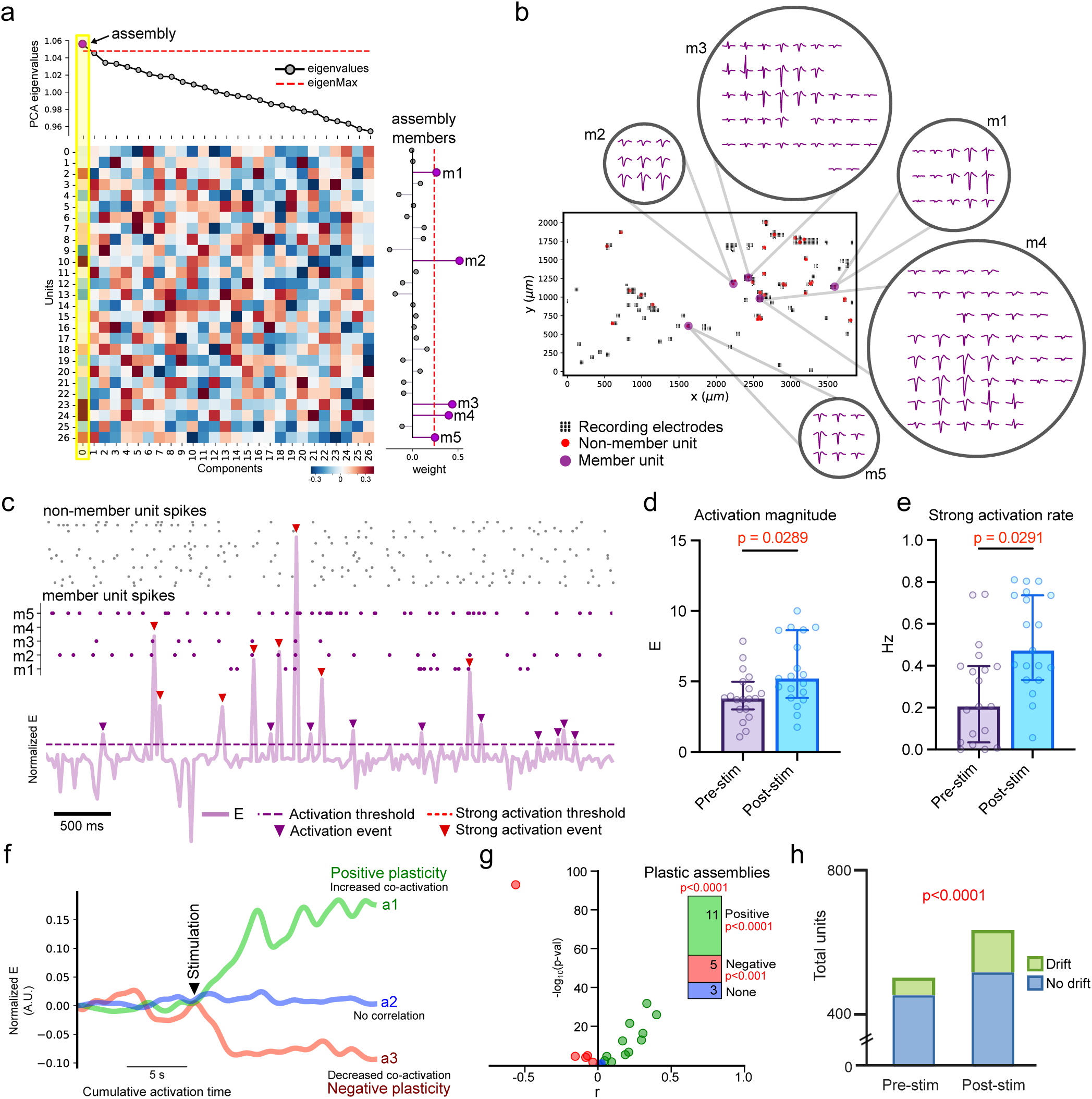
Cell assemblies are activated and strengthened after stimulation. **a**. Example of assembly identification with the combined PCA/ICA method. Heatmap shows the PCA components with associated eigenvalues in line plot at the top. Values for the identified assembly are highlighted in yellow. Right stem plot shows assembly members (m1-m5) passing the unit vector membership threshold derived from ICA results. **b.** Member unit locations and spike waveform footprints of assembly identified in (**a**) **c.** Magnitude of assembly activation strength *E* corresponds with co-activation of member neurons. **d.** Activation event magnitude increases after stimulation (paired t-test, t(n=19) = 2.375, p = 0.0289). **e.** The rate of strong activation events per second is higher after stimulation (paired t-test, t(n=19) = 2.372, p = 0.0291). **f**. Line graphs of activation strength over cumulative time are examples of three assemblies with different correlation (plasticity) profiles. Values shown are normalized activation strength *E* during activation events, with baseline set to zero and smoothed with a gaussian kernel. Tick indicates the transition from pre-stimulation to post-stimulation. **g.** Correlation between assembly event activation magnitude and the time of activation events is mostly positive. All comparisons are Spearman rank correlations. Inset shows proportion of assemblies with positive, negative, or insignificant Spearman’s r. Permutation tests on total number of assemblies, total positive and total negative assemblies ****p<0.0001, ***p<0.001. **h.** Neuron drift into and out of assemblies increases after stimulation (Fisher’s exact test, ****p<0.0001, odds ratio = 2.03).

We next examined the identity of assembly member neurons according to putative cell type cluster and cortical layer location. Assembly member neurons belonged to all seven cell type clusters without bias (**Supp Fig 2d**). There was no location preference when comparing member neurons in upper versus lower cortical layers (estimated as described in methods). However, member neurons were overrepresented in the 2000-2500um range of the MEA, which roughly coincides with cortical layer 4 (Fisher’s exact test, odds ratio = 1.93, p = 0.025; **Supp.** Fig 2e). These results demonstrate that assembly member neurons were diverse in cell type and location, with a preference for member neurons within the region of layer 4.

### Stimulation promotes local circuit remodeling through neural assembly activation and flexibility

To understand how stimulation affected the extracted assemblies, we calculated *E*, the strength of assembly member co-firing activity relative to population spiking activity (*E* is known as “expression” elsewhere in the literature [17], see methods; **Fig 2c**). In the post-stimulation period, the magnitude of *E* during activation events was higher than in pre-stimulation activation events (paired t-test, t(18) = 2.375, p = 0.0289; **Fig 2d**). In addition to this, we found that strong activation events occurred more frequently after stimulation (paired t-test, t(18) = 2.372, p = 0.0291; **Fig 2e**). These results indicate that stimulation promotes organized cell assembly activity rather than non-specific changes in the firing rate of individual neurons.

We further examined the relationship between stimulation and assembly activation by determining whether there was a correlation between activation event magnitude and time. A significant correlation between event magnitude and time represents an assembly activation profile that we refer to as either positive or negative “plasticity,” meaning that the connections driving the cofiring within an assembly significantly strengthened or weakened following stimulation (**Fig 2f**). We found a significant number of assemblies exhibiting a correlation between time and the magnitude of each activation event with 11 of 19 assemblies indicating a positive plasticity profile (**Fig 2g**). We implemented a shuffle control and confirmed that these correlations were significant beyond chance (**Supp.** Fig 2f).

As a separate analysis of local circuit plasticity, we next used previously established methods to test for assembly member flexibility, assayed by determining the number of neurons drifting in or out of assemblies specifically during periods of assembly activation [17]. We found a larger number of neurons exhibiting plastic changes after stimulation compared to baseline (assayed using the drift fraction, Fisher’s exact test, odds ratio = 2.03, p<0.0001; **Fig 2h**). We confirmed that the increase in drift fraction following stimulation was significantly greater than chance using a shuffle control procedure (permutation test, p<0.0001; **Supp.** Fig 2g). We note that since assembly activation events are calculated relative to overall spiking activity for putative member neurons, this result is not explicable due to non-specific firing rate changes. Overall, the evidence for increased assembly flexibility after stimulation both through new member neuron recruitment and enhanced assembly coactivation is consistent with the hypothesis that stimulation promotes plasticity in local neural circuits.

### Multiomic profiling of *ex vivo* stimulation in the human temporal cortex

To determine the molecular pathways affected by stimulating the human temporal cortex, we prepared multiomic snRNA-seq and snATAC-seq (snRNA/ATAC-seq) libraries of stimulated and non-stimulated temporal cortex slices from six subjects. After quality control, we retained a total of 50,067 nuclei with high quality snRNA/ATAC-seq data. Each nucleus had a median of 1,754 genes, 3,456 UMIs, and 5,957 ATAC peak fragments. We integrated the paired snRNA/ATAC-seq data, performed UMAP clustering and annotated clusters based on established marker genes and references datasets [23]. We detected 7 major cell classes including excitatory and inhibitory neurons, glial cells, and vascular cell populations, with similar contributions to cell types from all samples (**Fig 3a-b; Supp.** Fig 3). We next isolated and sub-clustered the excitatory neurons, inhibitory neurons, and non-neuronal cell types separately (**Fig 3c-d; Supp.** Fig 4). We further refined the annotation of the subclusters using a published human middle temporal gyrus reference dataset and made modifications to the naming conventions where necessary [23]. We detected 12 excitatory neuron subtypes and 16 inhibitory neuron subtypes.

**Figure 3.**
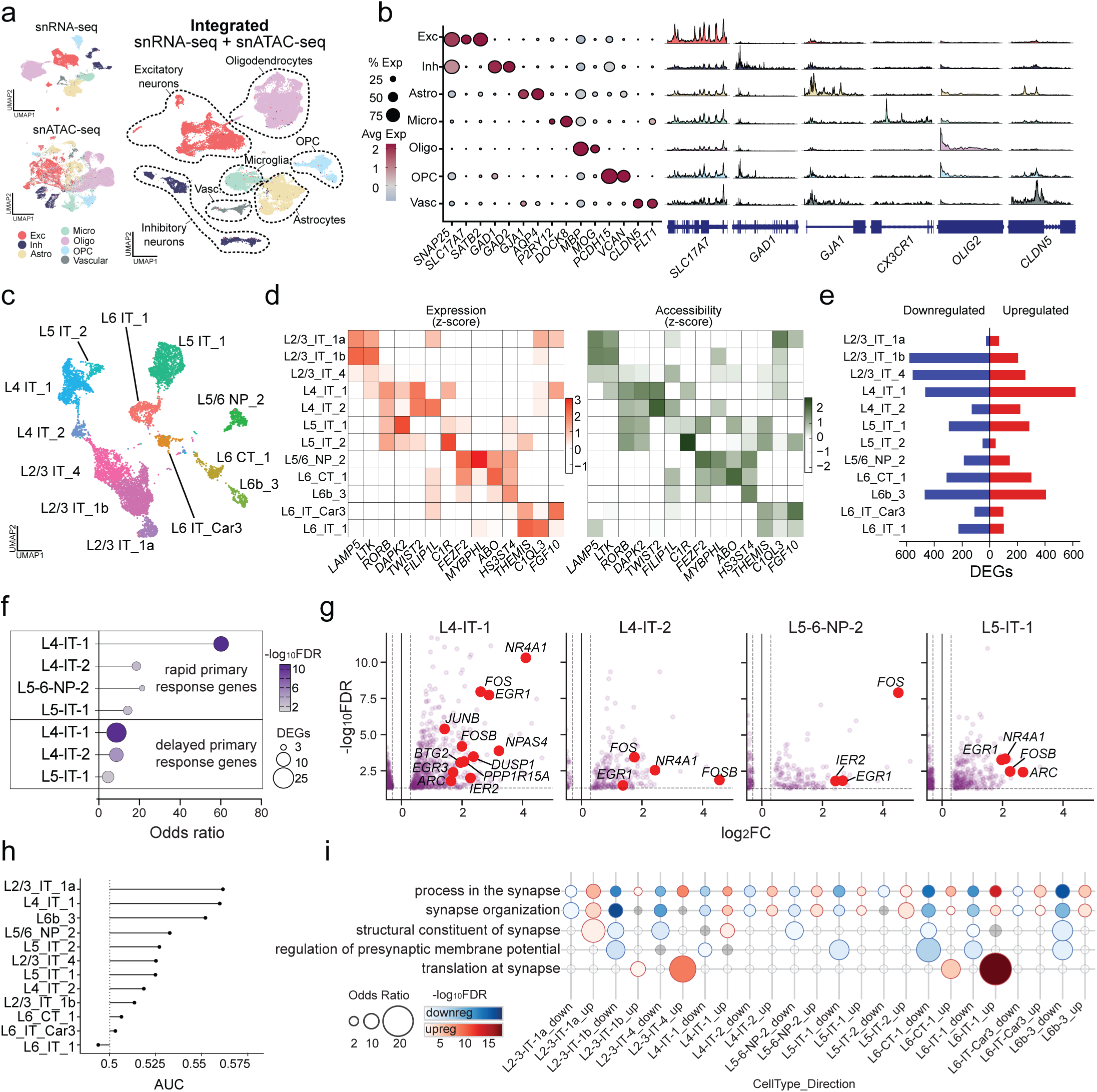
Stimulation induces differential expression of genes underlying activity and synaptic processes in excitatory neurons. a-b. Clustering of integrated single nucleus RNA and ATAC sequencing data (n = 50,067 nuclei from 12 samples [6 per condition]). **c-d.** Subclustered excitatory neurons. *n =* 7992. **e.** Number of downregulated and upregulated genes identified with pseudobulk differential gene expression analysis with DESeq2. **f.** Enrichment analysis for rapid and delayed primary response genes in the DEGs of each cell type. **g**. Differentially expressed rapid primary response genes are enriched in L4, L5, and L6 excitatory neuron types. **h.** Augur results point to L2/3-IT-1a and L4-IT-1 as the highest cell type priorities. **i.** Synaptic gene enrichment analysis using SynGO database reveals prominent enrichment of DEGs across multiple synaptic locations and processes. Upregulated and downregulated DEG lists from each list were analyzed separately.

### Specific excitatory neurons exhibit broad transcriptomic changes after stimulation

Pyramidal-like cells appeared to have the largest increase in firing rate after MEA stimulation. This effect was variable, though, with some cells also showing decreased or unchanged firing rate (**Fig 1g**). For this reason, we first asked whether specific excitatory neuron subtypes had altered gene expression upon stimulation. We carried out differential gene expression analysis and uncovered hundreds of differentially expressed genes (DEGs) in most excitatory neuron types (**Fig 3e; Supp. Table 3**). Compared to the other major cell classes, we saw the highest number of DEGs in excitatory neurons (**Supp.** Fig 5**; Supp. Table 3**). Layer 4 cells had the most DEGs, the majority of which were upregulated. Upper layer (L2/3) excitatory neuron subtypes also exhibited a large number of DEGs, mostly in the downregulated direction.

To determine whether stimulation led to differential ARG expression in a cell type-specific manner, we conducted an enrichment analysis for different classes of ARGs as defined previously by their activation time windows [10]. We found that rapid primary response genes (rPRGs) were enriched in the upregulated DEGs of four excitatory cell types: L4-IT-1, L4-IT-2, L5-IT-1, and L5/6-NP-2 (**Fig 3f-g**). Delayed PRGs (dPRGs) were significantly enriched in L4-IT-1, L4-IT-2, and L5-IT-1 cells as well (**Fig 3f**). As expected, secondary response genes (SRGs) were not enriched, which confirms that we are capturing a response specific to stimulation based on the timing of the experiments. Notably, of the 55 differentially expressed ARGs in L4-IT-1 cells, all but three were upregulated.

We next used Augur [24], an unbiased machine learning-based tool, to quantify and rank the transcriptomic response to stimulation in each excitatory neuron type. This cell type prioritization algorithm can identify meaningful but subtle biological responses to stimulation that may otherwise go unnoticed if using differential gene expression analysis alone. The top two candidate excitatory neuron types were L2/3-IT-1a and L4-IT-1 (**Fig 3h**). Compared to other excitatory subclusters, L2/3-IT-1a possessed the fewest number of DEGs and had few ARGs altered with stimulation. L2/3-IT-1a cells prominently expressed rPRGs like *FOS* and *ARC* at baseline and therefore our results may indicate a ceiling effect where stimulation is unable to further induce expression of these genes.

We sought to identify common molecular pathways outside of ARGs that are affected by stimulation in both L2/3-IT-1a and L4-IT-1 cells by identifying DEGs changing in the same direction in both cell types. We found 23 shared upregulated DEGs and 9 shared downregulated DEGs (**Supp. Table 4)**. Genes associated with the synapse (e.g., *HOMER1*) comprised 14 of the shared DEGs. Pathway enrichment analysis revealed significant enrichment for genes encoding members of receptor tyrosine kinase signaling cascades including a regulatory pathway of GABAergic signaling involving BDNF (*GABRG2, PIK3R1, NTRK2*). The common DEGs between the two cell types point to signaling pathways and specific gene targets like *HOMER1* that may be particularly important in the neuronal response to stimulation, but the larger magnitude of divergent characteristics demonstrates the unique impact stimulation has on each cell type. These findings underscore that stimulation not only activates rapid-response transcriptional programs in specific excitatory subpopulations but also reveals unique molecular signatures and pathways that distinguish their transcriptional responses.

### Differential expression of genes related to neural circuit modulation after stimulation

In addition to upregulated ARG expression, we hypothesized that stimulation would alter the expression of genes encoding synaptic proteins. Indeed, we found enrichment of synaptic genes in the DEGs of many excitatory neuron subclusters (**Fig 3i; Supp.** Fig 6**; Supp Table 5**). We did not see the same level of synaptic gene enrichment in inhibitory neurons, nor in most non-neuronal cell types (**Supp.** Fig 6**; Supp. Table 5**).

Genes associated with the pre-synaptic and post-synaptic compartments were highly represented in most excitatory subtypes, as were genes associated with synaptic signaling and organization. Terms uniquely enriched in upregulated DEGs were related to the synaptic cytoskeleton and regulation of the synaptic vesicle cycle. Terms unique to downregulated DEGs were related to axo-dendritic transport and the regulation of synaptic membrane potential. We also identified many differentially regulated voltage-gated ion channel genes (**Supp.** Fig 7), perhaps indicating that a homeostatic ion channel regulation process was occurring in response to increased cell depolarization [25, 26]. Taken together, these results show that stimulated excitatory neurons may be undergoing a synaptic remodeling process in support of local neural circuit connections.

### Establishing cell type-specific gene regulatory networks underlying stimulation responses

To explore how regulatory elements influence gene expression in excitatory neurons following stimulation, we constructed cell-type-specific maps of non-coding element-gene pairs under stimulated and control conditions. This was achieved by correlating chromatin accessibility peaks with gene expression levels in each condition using scMultiMap [27]. We identified 6,317 significant peak–gene links in excitatory neurons. Of these, 3,448 links were detected under stimulation and 2,869 under control, representing approximately a 1.20-fold increase in detected links during stimulation compared to control. This stimulation-associated increase in peak–gene linkage was consistently observed across multiple excitatory neuron subclasses, in particular L2/3 and L4 pyramidal neurons (**Supp.** Fig 8a). Additionally, we found many differentially accessible chromatin regions (DARs) in several excitatory neuron subtypes including L4 pyramidal cells (**Supp.** Fig 8b). These results reveal that stimulation amplifies the regulatory connectivity landscape in excitatory neurons, underscoring a dynamic, cell type-specific chromatin-to-gene coordination that likely underpins rapid transcriptional responsiveness following stimulation.

Building on this finding, we next sought to investigate the upstream drivers orchestrating these gene expression programs. Gene expression programs are typically organized by key transcription factors (TFs) that can enact multiple parallel signaling cascades driven by single inputs. These factors can be identified by combining gene expression readouts with TF motif enrichment within accessible chromatin regions. We leveraged our integrated snRNA-seq and snATAC-seq data to infer multifaceted gene regulatory networks (GRNs) using SCENIC+. This approach enabled us to reconstruct enhancer–driven GRNs at single-cell resolution, directly integrating chromatin accessibility with transcriptomic activity. (**Supp.** Fig 9a-b)

In the excitatory neuron GRN we identified 153 regulons consisting of 145 transcriptional activators, 8 repressors, and 115 unique primary TFs (**Supp. Table 6**). We determined the activity of each regulon at the level of individual cells using AUCell [28] to calculate the area under the curve (AUC) based on gene expression or accessibility of a regulon’s targets. We found that the cell type-specific regulon activity in excitatory neuron subtypes mirrored results from a study on the developing human neocortex [29]. For example, the SMAD3_+/+ regulon primarily operated in L2/3 excitatory neurons, while the NFIA_+/+ regulon was activated in L6 excitatory neurons (**Supp.** Fig 9c).

### Gene regulatory networks driven by ARGs are activated by stimulation in deep layer excitatory neurons

To understand the impact of stimulation on the GRN, we examined the differences in gene-based AUC scores between conditions and cell types. We found that stimulation led to significant activation of specific regulons in several excitatory neuron subtypes across the cortical layers (**Fig 4a-b; Supp. Table 7**). Conversely, regulons in interneuron subtypes appeared largely unaffected by stimulation (**Supp.** Fig 10**; Supp. Table 7**). This is congruent with the limited differential gene expression response we observed in inhibitory neurons (**Supp.** Fig 5**; Supp. Table 3**) in addition to the relatively small firing rate changes we recorded in putative interneurons (**Fig 1f-g**).

**Figure 4.**
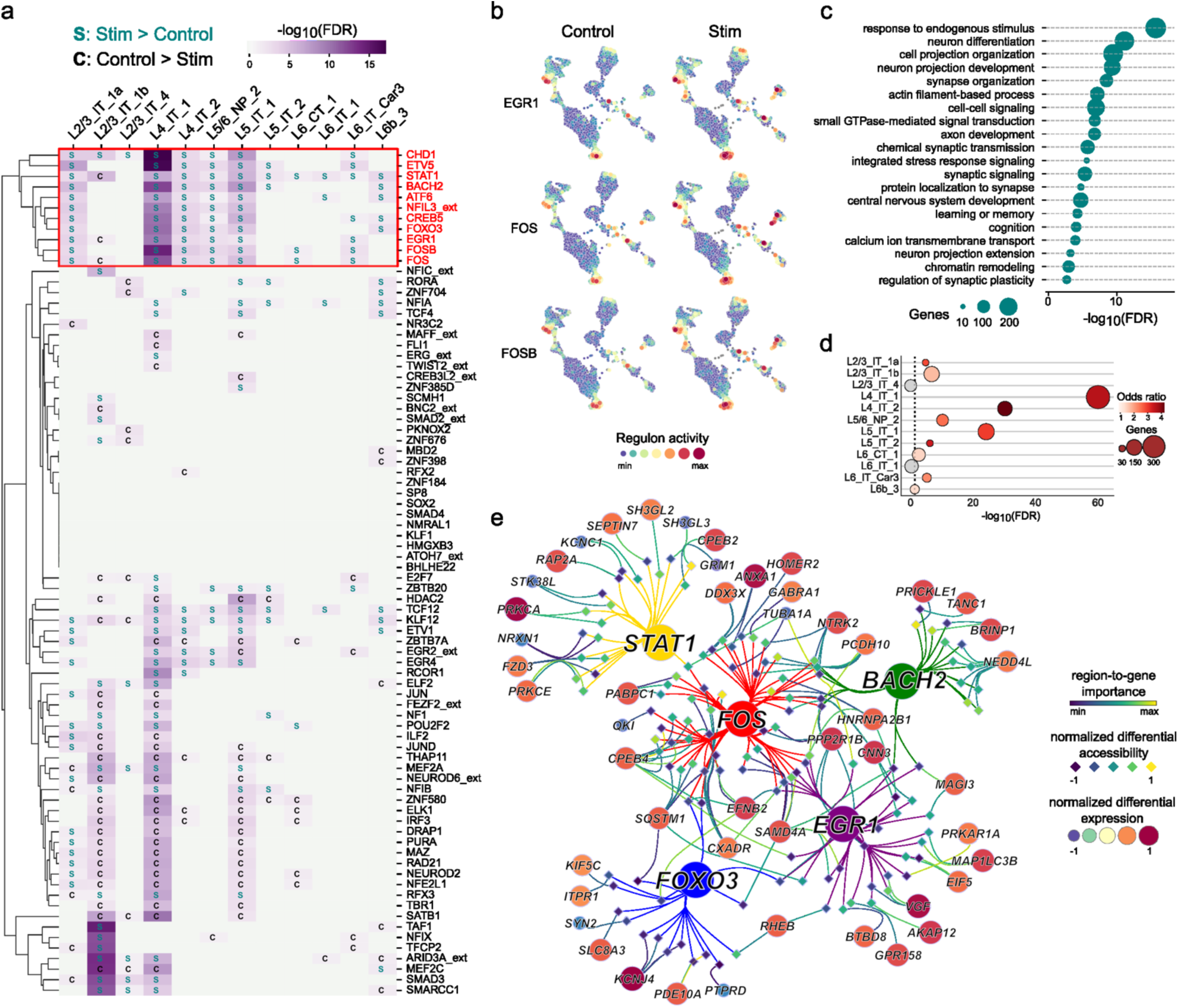
Stimulation engages activity-dependent gene regulatory networks in deep layer excitatory neurons. **a**. Comparison of regulon activity in control and stimulated cells of each type. A network of several regulons (red box) is prominently activated in select excitatory neuron subtypes. Primary transcription factors in this network include immediate early genes. **b.** Feature plots of normalized activity scores for EGR1, FOS, and FOSB regulons in control and stimulated excitatory neurons. **c.** Gene ontology enrichment analysis of all target genes in the highlighted network in (**a**). **d.** DEGs of several cell types are significantly enriched in the target genes regulons in the highlighted network in (**a**). **e.** Select primary transcription factors (central circles), target regions (diamonds), and target genes (peripheral circles) of the highlighted network. Each target gene is related to the synapse and is also a DEG in L4-IT-1 cells. Color and size of target gene symbols represent differential expression in L4-IT-1 cells. Edges between transcription factors and regions are colored according to transcription factor. Width, color, and transparency of region to gene edges correspond to the region-to-gene importance scores.

One group of regulons was consistently activated in the four deep layer cell types with enrichment of differentially upregulated ARGs and also in the upper layer neuron subcluster flagged by Augur prioritization (**Fig 3f-h; Supp. Table 7**). This network was most notably activated in L4-IT-1 cells. Primary TFs active in this network included the ARGs *FOS* and *EGR1* as well as other TFs known to participate in the response to cellular activation and neuronal plasticity like *CHD1* and *ETV5* [30, 31]. Target genes were involved in synaptic signaling and organization, neurodevelopment, and learning and memory (**Fig 4c**). The target genes were also enriched for DEGs of several cell types, and to the greatest degree in L4-IT-1 DEGs (**Fig 4d**). Many of the target genes that overlapped with L4-IT-1 DEGs were upregulated synaptic genes (**Fig 4e**). For example, *HOMER2, SAMD4A,* and *PRICKLE1* are upregulated target genes that are critical for glutamatergic synapse assembly [32–34]. Additionally, *AKAP12* and *SQSTM1* are upregulated target genes that are essential for synaptic plasticity [35–37]. Together, these results indicate that stimulation dynamically reshapes the genomic landscape to support cell type-specific transcriptional responses that may underly neural circuit modulation.

### Non-neuronal participation in the response to stimulation

An advantage of applying stimulation to an organotypic slice preparation, in contrast to a neuronal cell culture method, is that we are able to examine the response of non-neuronal cell types as well. We identified two astrocyte subtypes in our data (**Fig 5a-b**). Both subtypes reacted to stimulation with differential expression of genes related to synapses, cell excitation, and gap junctions (**Fig 5c-d; Supp. Table 3**). Augur analysis tagged the Astro_4 subtype as high priority due to the effect of stimulation on its transcriptomic profile (**Fig 5e**). Genes encoding gap junction proteins were upregulated in Astro_2 cells (*CDH19, JAM3*) and Astro_4 cells (*GJC*) (**Fig 5d; Supp. Table 3**), suggesting that connections between astrocytes are being strengthened. Stimulating a small group of astrocytes can cause waves of activation across large populations of astrocytes which can ultimately enhance or inhibit excitatory synaptic signaling depending on the context [38]. Astrocytes also release vesicles containing neurotransmitters using a similar molecular pathway to synaptic vesicles [39, 40]. *SV2C* and *PACSIN2* were upregulated in astrocytes (**Fig 5d; Supp. Table 3**), both of which are involved in the regulation of synaptic vesicle exocytosis [41, 42]. Also upregulated is *PTX3* (**Fig 5d; Supp. Table 3**) which encodes a soluble factor that is secreted by astrocytes to promote the formation of excitatory synapses [43]. Altogether, the upregulation of these genes supports a role for astrocytes participating in the coordinated regulation of synaptic modulation and excitability.

**Figure 5.**
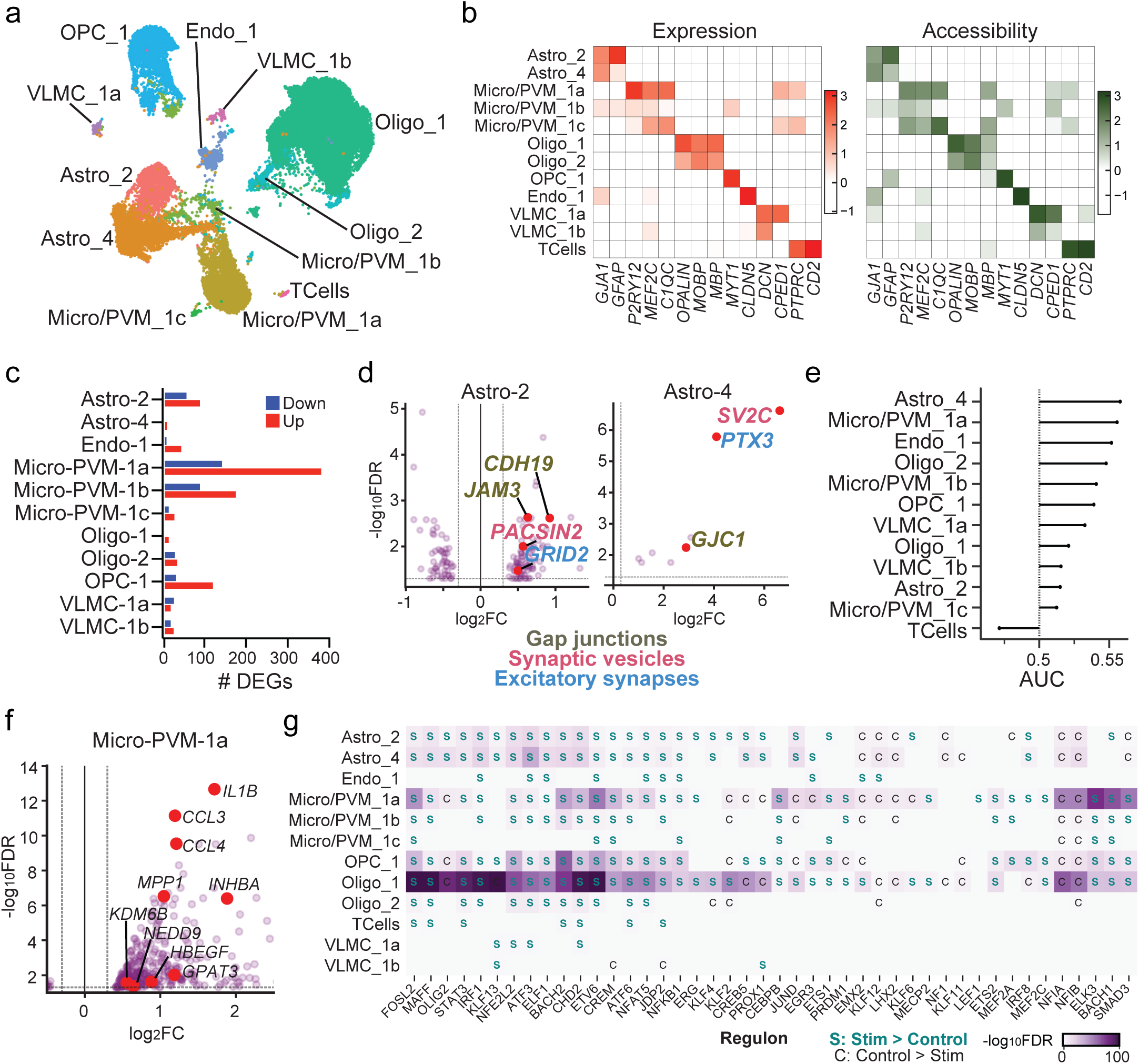
Stimulation differentially activates gene regulatory networks in non-neuron subtypes. **a**. Non-neuron subclusters. **b**. z-scored expression and chromatin accessibility of subcluster marker genes. **c.** Number of downregulated and upregulated genes identified with pseudobulk differential gene expression analysis with DESeq2. **d.** Select astrocyte DEGs related to gap junctions, synaptic vesicles, and excitatory synapses. **e.** Augur results point to Astro_4 cells as the highest non-neuron type priority. **f.** Highlighted microglial DEGs overlap with results from a study on *in vivo* stimulation of the human temporal cortex [45]. **g.** Differential activation of select regulons in specific non-neuron subtypes.

Microglia are known to participate in synaptic pruning [44], but their role in neuromodulation mediated circuit remodeling is unknown. We found several hundred microglial DEGs, including dPRGs like *NR4A3* and synaptic genes like *HOMER1.* A subset of these DEGs overlapped with microglial DEGs identified in a recent manuscript on stimulation of the human cortex *in vivo* (**Fig 5f**) [45]. These included *CCL3, CCL4,* and *IL1B*, all encoding cytokines. This overlap indicates that our stimulation paradigm is affecting microglia in a similar manner as *in vivo* stimulation, which is evidence that we are not causing abnormal inflammatory reactions.

Interestingly, *SIRPA* was upregulated in Micro-PVM-1a cells. *SIRPA* encodes an important anti-phagocytic signaling receptor that protects against excessive synaptic pruning, and its loss is accompanied by learning and memory deficits [46]. Furthermore, post-mortem studies of human Alzheimer’s disease brains uncovered a correlation between disease advancement and downregulation of *SIRPA* [46]. Microglial activation is seen following the insertion of depth electrodes in the human brain, which contributes to an inflammatory profile that can be deleterious in the acute and chronic periods [47–49]. However, our results indicate that stimulation may be used to promote a more senescent microglial phenotype associated with positive effects at the synapse.

### Stimulation alters glial gene regulatory networks

Next, we aimed to understand the effect of stimulation on GRNs in non-neurons. We identified 162 regulons controlled by 134 primary transcription factors in non-neuronal populations (**Supp. Table 6**). Astrocytes, oligodendrocytes, OPCs, and microglia showed activation of many regulons after stimulation (**Fig 5g**). Regulons activated in astrocytes included KLF4 and FOSL2 (**Supp. Table 7**). Other activated astrocyte regulons like CREB5, KLF13, and PROX1 targeted genes involved in cell-cell adhesion, the response to calcium, and ion transport at cell appendages (**Supp Fig 10a-c**). We also found that stimulation altered regulon activity in oligodendrocytes and their precursor cells (**Fig 5g**). Together, these results support the model that stimulation and/or neural activity excites astrocytes through calcium-mediated mechanisms that can propagate across gap junctions and modulate synapses. Furthermore, these results indicate that oligodendrocytes may be working in tandem with other non-neurons to modulate neural excitability in a manner that strengthens cell circuit connections.

## Discussion

Here we demonstrate that brain stimulation modulates local cortical circuits and engages cell type-specific gene regulatory networks. This is the first time that gamma assemblies have been identified in the human temporal cortex *ex vivo*. *In vivo,* gamma assemblies supporting human memory performance have been found to exhibit enhanced activation and drift capabilities [17]. We saw similar phenomena in the stimulated temporal cortex *ex vivo*, suggesting that stimulation alters cell assemblies in a manner that may support cognitive processes *in vivo*. Moreover, we uncovered a stimulation-sensitive GRN in excitatory neurons containing target genes related to cellular activation, synaptic plasticity, and learning and memory. The GRN was prominently activated in specific excitatory neuron types in L4 and L5. This, along with the relative lack of changes seen in the inhibitory neuron GRN, mirrors our electrophysiology results in which putative pyramidal cells exhibited the highest increase in firing rate after stimulation. Additionally, we found that assembly member neurons were more likely to be located in the region corresponding to cortical layer 4. Together, these results suggest that excitatory neurons throughout the deep layers of the cortex are the primary targets of this stimulation paradigm. Previous investigations of gene expression changes in rodents subjected to DBS have suggested that stimulation enhances transcription of genes associated with neuronal survival and synaptic plasticity [9]. Our data are consistent with these existing models and also add important information about stimulation in cell type- and human-specific contexts.

We identified neuronal assemblies on time scales defined by a cycle of low gamma oscillations based on previous work in both rodents and humans [14, 17, 50]. This time scale facilitates long term potentiation and organization of individual firing through hierarchical interactions of theta and gamma oscillations [16, 51]. We certainly acknowledge that cofiring on other time scales is possible, and that further characterization of e*x vivo* physiology and response to stimulation will require explication of optimal frequencies that modulate cofiring. We note however that the identification of assemblies, especially with the computationally rigorous methods we employ [22], provides remarkable correspondence with *in vivo* experimentation [17]. Alternative mechanisms to assay stimulation-related circuit plasticity represent a goal of subsequent experimentation. This may include methods such as optogenetics to specifically activate neurons of interest (e.g., layer 4 pyramidal cells) in order to gauge their direct role in cell assembly modulation. The feasibility of utilizing optogenetics in human organotypic slices in conjunction with MEA recordings was recently demonstrated [52]. Similar approaches have been used to define and enhance the cell type-specific mechanisms of DBS for motor disorders and therefore may be translated to our paradigm as well [53, 54].

The tissue preparations we employed here, which preserve cortical layers, provide a unique and unparalleled basis for understanding the effect of stimulation on local circuits. However, it does preclude the adoption of stimulation parameters that are more directly analogous to those employed *in vivo*. We explicitly sought parameters that 1) would not specifically elicit local potentiation or depression [55–57]], as we believed this could skew our findings; 2) had been proven not to elicit tissue injury [58]; and 3) utilized analogous charge transmission to the tissue as in DBS paradigms [59]. The significant and specific impact of stimulation on neurophysiology supports the relevance of our findings more widely, although we fully acknowledge that additional testing with several stimulation waveforms will be necessary to fully understand our results and place them in the context of clinical brain stimulation.

The role of gene expression changes in glia in response to stimulation complements our principal results identifying alterations in neuron populations. Our results align with previous studies on ARG induction in astrocytes [60] in addition to recent evidence of stimulation inducing differential gene expression in glia in human temporal cortex *in vivo* [45]. There is increasing evidence that astrocytes facilitate the therapeutic effects of stimulation via regulation of synaptic plasticity and neuronal excitability [61]. Other emerging research shows that oligodendrocytes can sense and respond to neuronal activation to enable the dynamic regulation of axonal excitability and conduction via myelination, which may contribute to successful learning and memory processes [62–64]. There is evidence that oligodendrocytes enhance myelination in response to neuronal activity in part due to adenosine signaling [63, 64]. Adenosine appears to be an important mediator of neuromodulation therapies [65, 66], which may be due to its effects on oligodendrocytes in addition to networks of astrocytes and neurons.

Ultimately, this work has defined specific gene networks underlying stimulation-induced neural circuit plasticity. Future work should incorporate additional samples and stimulation paradigms, including *ex vivo* chronic stimulation and *in vivo* stimulation of the human temporal cortex. Building on the present study, these additional strategies will uncover translational cell type-specific mechanisms of neuromodulation in the human temporal cortex, potentially resulting in the development of effective therapies for cognitive disorders.

## Methods

### Tissue collection and processing for *ex vivo* experiments

All tissue was donated by fully consented patients undergoing surgical treatment for epilepsy under the IRB protocol STU 092014-026. We did not use tissue directly containing seizure foci or other pathologies like tumors or cortical dysplasia. Brain tissue collection, dissection, and slicing were completed in ice cold, carbogenated N-methyl D-glucamine (NMDG) aCSF (92mM NMDG, 2.5mM KCl, 1.25mM NaH2PO4, 30mM NaHCO3, 20mM HEPES, 25mM D-Glucose, 2mM Thiourea, 5mM Na ascorbate, 3mM Na pyruvate, 0.5mM CaCl2, 10mM MgSO4, pH 7.3, 300-310 mOsm) [67].

Brain samples were obtained at the time of temporal lobectomy operations. As is standard for these procedures, the anterior temporal cortex was resected *en bloc*. Then, the neurosurgeon dissected a small portion (∼3 cm^3^) and placed it into sterile, ice-cold NMDG aCSF. We promptly (<10 minutes) transported the tissue sample on ice to a sterile biosafety cabinet for further processing (**Fig. 1a**). We dissected away large blood vessels, pia mater, and areas exposed to electrocautery during surgery. We then prepared a block of tissue (∼1 cm^3^) with a clear cross-section of the cortex from the pial surface to the white matter. The tissue block was then embedded in low-melting point agarose (3% in sterile PBS, held at 37C), then sectioned using a Compresstome (VF-510-0z, Precisionary Instruments) at a thickness of 250 uM. After sectioning, the slices rested for a period of 12 minutes in warmed, carbogenated NMDG aCSF. This resting period was followed immediately by preparation for culture or recordings.

### Human organotypic slice culture

We based our method and organotypic slice culture medium (OSCM) on established protocols that have previously been used to culture human cortex slices for long periods of time [68, 69]. We prepared fresh OSCM and high HEPES OSCM (H-OSCM) prior to each tissue collection as described in [69]. After resting in warm NMDG aCSF, we transferred the slices to permeable cell culture membranes (Millipore cat no. PICM03050) in 6-well plates containing 1 mL H-OSCM per well. After recovering for one hour in a tissue culture incubator (37C, 5% CO2, 100% humidity), we transferred the inserts to new 6-well plates containing 1 mL OSCM per well. We supplemented the OSCM with Primocin (2 uL/mL; Invivogen cat no. ant-pm-05) for the duration of the culture period. We completed a 50% media change daily. The day of resection was defined as 0 days *in vitro* (DIV0), and each day thereafter as DIV1, DIV2, and so on.

### HD-MEA recordings

We used the MaxWell Biosystems MaxOne device for recordings and stimulation. We first transferred from the NMDG aCSF bath or from their culture membranes to warm, carbogenated normal aCSF (123mM NaCl, 4mM KCl, 25mM NaHCO3, 1.2mM NaH2PO4, 10mM D-Glucose, 1mM CaCl2, 1mM MgSO4, pH 7.3-7.4, 290-310 mOsm) [70] to equilibrate for at least 45 minutes. We prepared each MEA chip by soaking the electrode surface in MilliQ water overnight. Prior to placing the slice, we removed the water and rinsed the chip two times with normal aCSF. We transferred the slice to the chip using a glass transfer pipette. We carefully manipulated the slice using a paintbrush to position it over the electrode array, always with the white matter facing the same direction for every slice (**Fig. 1a**). After lowering the tissue holder into place, we began a perfusion of warm, carbogenated aCSF at a speed of 1 to 2 mL per minute.

We used preset recording assays defined in the MaxLive recording software. Every recording was executed with a sampling rate of 20 KHz, hardware high pass filter of 1 Hz, and gain of 512. First, we completed an “activity assay” which records activity at multiple electrode configurations covering the entire electrode array. The electrodes with the highest amount of activity in the initial activity assay are included in the electrode configuration that is used in the subsequent pre-stimulation “network assay”. The network assay is a continuous recording at a defined electrode configuration. After stimulation, the network assay is repeated with the same electrode configuration that was used pre-stimulation. These two network assays were concatenated and used to identify single units and assemblies in downstream analyses.

### Stimulation

We designed a custom stimulation program in Python using the MaxWell Biosystems API. The fifteen-minute stimulation period was composed of several cycles, each consisting of five bursts of five bipolar, positive-leading square pulses (5 s inter-burst interval, 50 ms inter-pulse interval, 200 us phase length, 300 mV; **Supp.** Fig 1a). To avoid specifically stimulating any one region of the slice, a new group of 32 stimulation electrodes was configured after each stimulation cycle. Parameters were chosen based on previous *in vitro* studies demonstrating tissue safety, lack of inducing long-term potentiation or depression, and for similarity to those used *in vivo* for clinical applications [55, 56, 58].

### Scan-level activity analysis

We completed 37 stimulation experiments consisting of the recordings and stimulation periods described above. To identify action potentials, we used a spike detection threshold of 5 root mean square over the background voltage signal. We determined the average 90^th^ percentile firing rate (FR90) and the average spike amplitude recorded across all electrodes in the pre- and post-stimulation scans. For the scan-level activity analyses, we only included experiments with >0.1% active electrodes in the pre- and post-stimulation scans, where an active electrode is defined as having an average FR of at least 0.1 Hz and an average spike amplitude of at least 10 uV. Of the 37 experiments completed, 29 passed this inclusion threshold. We used a paired-sample Wilcoxon signed-rank test to compare FR90 and a paired-sample t-test to compare spike amplitudes of the pre- and post-stimulation scans (*n* = 29).

### Spike sorting

We used the algorithm Kilosort (v2) [71] and Spike Interface (v0.101.0) [72] for spike sorting and quality control curation. We used a combination of automated and manual curation to confirm the quality of the identified units [73]. We excluded units with signal to noise ratios less than 2, inter-spike interval violations greater than 3%, and firing rates less than 0.1 Hz. In all single unit and assembly analyses, we focused only on units with negative peaks, as positive peaks are thought to represent axonal spikes [74]. We implemented rigorous spike sorting quality control procedures such as automatic curation based on signal to noise ratios, interspike interval violations, and minimum firing rate, as well as manual inspection of the location, waveforms, and autocorrelogram of each unit. Thus, we ensured that our analysis included well-isolated neurons rather than multiunit activity.

### Single unit classification – location

Cortical layer 4 is roughly 1500 uM away from the pial surface of the human temporal cortex [75]. We consistently positioned each slice so that the pial surface would align with the same edge of the MEA (Fig 1a). Thus, we defined the chip area within 1500 uM of the pial edge as the upper layers (L1-3) and anything below that as the lower layers (L4-6). We determined the location of individual sorted units by identifying the location of the electrode recording the largest spike amplitude for each unit. We worked under the supposition that the largest spike amplitude corresponds with the recording electrode closest to the cell body or axon initial segment.

### Single unit classification – putative cell type

We used the Python package WaveMAP to further classify sorted units into putative cell types [76, 77]. WaveMAP uses Louvain clustering and uniform manifold approximation and projection (UMAP) to group cells based on the shape of their waveforms. We extracted up to 500 spike waveforms from each unit at the electrode exhibiting the largest spike amplitude. The waveform traces consisted of 6 ms centered around the peak of the spike. We then averaged all waveforms for each unit, aligned all averaged unit waveforms at their peaks, and trimmed the traces to 1.5 ms before and 2 ms after the peak. We normalized the aligned, trimmed waveforms to fall between -1 and 1 arbitrary units as the final processing step before running the WaveMAP algorithm. We identified the best clustering resolution and number of neighbors by examining the modularity scores and number of clusters identified with a combination of the two parameters as described in [77]. Ultimately, we used a resolution of 1.5 and 20 neighbors to split the 494 units into 7 clusters.

### Single unit activity analysis

We calculated the firing rate (*FR*) for each unit:

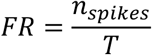

Where *n_spikes_* is the number of spikes occurring during recording time *T* in seconds. We calculated the change in firing rate (Δ*FR*) as:

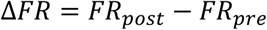

Where *FR_post_* is the firing rate for the post stimulation recording and *FR_pre_* is the baseline firing rate in the pre-stimulation recording. We used a paired-sample Wilcoxon signed-rank test to compare *FR_pre_* and *FR_post_*.

We also calculated the “modulation factor” (MF) [54] as a measure of firing rate change:

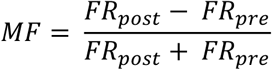

MF exists on a scale from -1 to 1, where -1 indicates that a neuron was completely silent in the post-stimulation period, 0 means no change in firing rate, and 1 indicates that a neuron was completely silent in the pre-stimulation period.

### Assembly identification

We identified assemblies using a two-step framework [22] that has been applied to intracranial recordings from rodents [50] and humans [17] to identify assemblies that participate in learning and memory. We searched for assemblies in concatenated recordings that had more than one unit with a mean firing rate greater than 0.5 Hz. This firing rate threshold is necessary to avoid skewing the data toward extreme values, which can lead to spurious PCA eigenvalues [17, 78, 79]. For each concatenated recording, we calculated the *z-*scored spike count of individual units in 25 ms non-overlapping time bins to generate a neuron-by-time bin spike rate matrix with a null mean and unitary variance.

We then performed a principal component analysis (*sklearn.decomposition.PCA()*) on the spike rate matrix. The resultant principal component by neuron matrix (*pca_results*.*components_*) is accompanied by a list of eigenvalues (*pca_results*.*explained_variance_*) of length equal to the number of components. Any principal component with an associated eigenvalue exceeding the upper bounds defined by the Marčenko-Pastur law [80]:

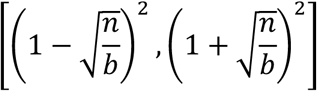

where *n* is the number of recorded neurons and *b* is the total number of time bins, indicates the presence of a significant co-firing relationship between neurons [79]. We used this eigenvalue threshold to isolate only the significant components into the neuron-by-assembly matrix 𝑃_sign_. We used a subsequent independent component analysis (*sklearn.decomposition.FastICA(max_iter=1000, random_state=1, whiten=’unit-variance’*)) to confirm the presence of assemblies and determine their member neuron compositions. Prior to running the ICA, we projected the rate matrix onto the principal components with significant eigenvalues:

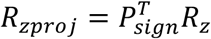

where *R_zproj_* is the projected spike rate matrix, 𝑃*_sign_^T^* is the transpose of the matrix of significant components for each neuron, and *R_z_* is the original *z*-scored spike rate matrix. We applied the ICA algorithm to *R_zproj_* to solve for the unmixing matrix, *U* (*ica_results.components_)*, which we then used rotate the PCA-derived assemblies to generate a neuron-by-assembly matrix of weight vectors, *W*:

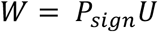

The weight vector represents the degree to which each neuron contributes to an assembly. To identify member neurons (i.e., neurons that significantly contribute to an assembly’s activity), we converted the weight vectors to scaled unit vectors, *V*, with the sign set so that the vector with the largest absolute value was positive [50].

Neurons are assembly members if they have a unit vector greater than a membership threshold defined as one standard deviation plus the mean of that assembly’s *V.* Any assembly with fewer than two member neurons is not considered a true assembly.

We employed a shuffle control to confirm that the assemblies we identified were not due to statistical chance. We randomly shuffled the spike times of each neuron (with inter-spike intervals preserved [81]), then reconstructed the firing rate matrix and applied the assembly identification analysis. We repeated this process 1000 times to generate a null distribution which we then compared to the number observed using a permutation test. [17]

### Assembly activation

To track the activity of each assembly over the length of the recordings, we calculated the assembly activation strength:

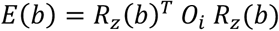

where 𝐸(𝑏) is the activation strength of assembly *i* at time bin 𝑏, 𝑅_z_(𝑏)*^T^* is the transpose of the normalized spike rate matrix column corresponding to bin 𝑏, and 𝑂*_i_* is the outer product of the unit vector of assembly *i*. The magnitude of 𝐸 depends on the number of active assembly members and the instantaneous firing rate of the members. Because we set the diagonal values of 𝑂*_i_* to zero, 𝐸 is not driven by the activity of a single member neuron and instead relies on member co-activation. We considered an assembly to be active in any time bin where 𝐸 exceeded the 95^th^ percentile. We observed the presence of strong activation events where 𝐸 surpassed the 99^th^ percentile and defined assembly activation rate as the number of strong activation events per second. Note that *E* is known as “expression strength” elsewhere in the literature [17, 50]. We instead use the term “activation strength” to avoid confusion with gene expression within this study.

### Assembly member drift and plasticity

Variability in membership after an experience is a feature of flexible assemblies that participate in memory updating. The drift fraction is a measurement of the number of neurons joining (drifting in) and leaving (drifting out) an assembly relative to the total number of neurons in a recording [17]. To determine whether neurons were drifting into or out of an assembly, we executed a Spearman rank correlation on a neuron’s z-scored firing rates during assembly activation events versus the time of those events. A member neuron with a significant negative correlation (p<0.05, r<0) is drifting out while a non-member neuron with a significant positive correlation (p<0.05, r>0) is drifting in. We calculated overall drift fractions [17] by summing the total number of drifting neurons and the total number of non-drifting neurons for each assembly before and after stimulation, then used a Fisher’s exact test to determine whether stimulation had a significant impact on the number of drifting neurons. We used a similar method to interrogate assembly plasticity, which we defined as the Spearman rank correlation between assembly activation strength (*E*) and activation event time across the entire concatenated scan. Assemblies with a positive correlation were strengthened by stimulation, indicating a positive plasticity profile among the assembly members. Assemblies with a negative correlation were oppositely affected.

To control for the possibility that the correlations we saw were due to chance, we implemented a shuffle control procedure wherein our observed value of interest was tested against a null distribution using a permutation test [17]. For drift fraction, we shuffled the order of activation events and calculated Spearman’s ρ for each individual neuron. We generated a null distribution of pre- and post-stimulation drift fractions by summing the total number of drifting and non-drifting units identified across all assemblies for each of the 1000 shuffles. We also compared the distribution of differences between the pre- and post-stimulation drift fractions to show that the increased drift after stimulation was not due to chance. For assembly plasticity, we generated a null distribution of Spearman’s ρ values from 1000 shuffled activation event arrays. For each assembly, we tested whether the observed Spearman’s ρ was significantly different from the null distribution. We also determined the total number of significant correlations identified in each iteration across all 19 assemblies and compared that null distribution to the number of observed significant correlations.

### Multiome sample preparation and nuclei isolation

Slices undergoing the MEA stimulation protocol on DIV0 were flash frozen immediately after the final recording was complete (∼15 minutes after stimulation). Control slices rested in normal aCSF for at least one hour before being flash frozen. We chose to include only DIV0 slices in the multiome experiments to prevent confounding gene expression changes that may occur during the early culture period [82]. The tissue slices are very small, weighing approximately 5 mg each. To have at least 20 mg of tissue for the multiome assay, we combined 4 slices originating from the same patient and condition per nuclei preparation.

The nuclei isolation protocol we used is an adaptation of demonstrated protocols for multiome complex sample preparation (10x Genomics, Demonstrated Protocol CG000375) and for snRNA-seq of human hippocampal tissue [83, 84]. We prepared all buffers as detailed in Protocol CG000375. We followed the lysis protocol described in [83] using a Dounce homogenizer for the initial tissue homogenization step, after which we incubated the sample in lysis buffer for 5 minutes on ice then spun down the samples at 500 x g in a tabletop microcentrifuge at 4C. We repeated the lysis and spin for a total lysis time of 10 minutes. We washed the pellet by soaking it in nuclei suspension buffer for 5 minutes on ice. We then resuspended the pellet, spun it down (5min,500xg, 4C), and removed most supernatant. We resuspended the pellet in nuclei suspension buffer and passed the sample through a pipette filter tip with 40 uM pores (Flowmi, BAH136800040) into a new tube.

Instead of doing the nuclei sorting procedure from Protocol CG000375, we used a sucrose gradient to clean up cellular debris from the nuclei samples. We followed the sucrose gradient protocol detailed in [83] using a tabletop microcentrifuge at 4C set to 13000 x g for 45 minutes. After the sucrose gradient, we washed the pellet in nuclei suspension buffer, spun it down, then resuspended in 100-200 uL nuclei suspension buffer. We combined 2 uL of sample with 18 uL Trypan blue and manually counted the nuclei. We then adjusted the volume of nuclei to equal 3,500 nuclei per uL, spinning and resuspending in the proper volume if necessary. We then completed the final nuclei permeabilization steps in Protocol CG000375 and proceeded to the multiome assay using 5 uL concentrated nuclei stock.

### Multiome data processing

Raw single-nucleus multiome sequencing data were analyzed using the Cell Ranger ARC v2.0.2 pipeline (10X Genomics). The Cell Ranger-ARC count pipeline was used for cell barcode calling, read alignment and quality check using the human reference genome (hg38, gencode v42) according to the protocols described by 10x Genomics. Nuclear fractions that quantify the amount of RNA in a droplet that originated from unspliced pre-mRNA were calculated for all cell barcodes in the sample BAM file using DropletQC (v.0.0.0.9000) [85]. Gene expression libraries in this project showed a high fraction of reads in nuclei, indicating high RNA content in called nuclei and low levels of ambient RNA detected. Gene expression and ATAC assays were separated using the Python package Scanpy (v1.9.6). For RNA data, we used Cellbender (v0.2.0) [86] to detect and adjust for ambient RNA contamination. Initial downstream analysis was performed separately.

For snTAC-seq data analysis, open chromatin region peaks were identified on individual samples using MACS2 (v.2.2.7) [87]. Peaks falling in the ENCODE blacklisted regions were excluded [88]. We selected peaks that were detected in more than ten cells. All samples were merged resulting in 250,810 processed ATAC peaks. For downstream analysis we retained the top 20% of consensus peaks across all nuclei. Latent Semantic Indexing (LSI) transform workflow was used which involves Term Frequency-Inverse Document Frequency (TF-IDF) for normalizing and Singular Value Decomposition (SVD) for dimensional reduction using the R package Signac (v.1.10.0) [89]. Peaks were annotated using ChIPseeker (v1.42.1) [90] based on the TxDb.Hsapiens.UCSC.hg38.knownGene annotation data available on Bioconductor. A transcription start site (TSS) window of +/-3kb was used for each gene, and genomic regions were automatically assigned by ChIPseeker. We created a gene activity matrix inferred from ATAC-seq by using the GeneActivity function in Signac, which assesses chromatin accessibility at gene body and promoter regions.

For snRNA-seq data analysis, downstream analysis was performed in R with Seurat (v4.3.0) [91] and customized R scripts. Data from different conditions (Control and Stimulation) were merged into a unique single Seurat object. Nuclear fraction score (described above) was added to Seurat metadata. Normalization and data scaling were performed using SCTransform v2 (v.0.4.1) [87] in Seurat.

Data integration of the two modalities (RNA and ATAC) was performed in R using Seurat and the following criteria: nuclei with > 250 genes, < 25,000 UMIs, < 5% mitochondrial transcripts, < 180,000 ncount_ATAC, < 2 nucleosome signal, > 1 TSS enrichment and > 0.5 nuclear fraction were retained for downstream analysis. Doublets were identified using scDblFinder (v1.12.0) [92] with default parameters. Batch correction was performed using Harmony (v0.1.1) [93]. Weighted nearest neighbor (WNN) graph was constructed according to the integrated dimensional reductions from two modalities with Seurat using 30 principal components and 50 LSI components. The resulting nearest-neighbor graph was used to perform UMAP embedding and clustering using the SLM algorithm [94]. We identified 26 final distinct clusters, and we identified cluster markers using a fast Wilcoxon rank sum test followed by auROC analysis developed in presto (v1.0.0). Initial cell annotation was done by manual curation of known canonical markers for excitatory neurons, inhibitory neurons, and non-neuronal cells. In excitatory neurons, clusters with low UMI and low number of genes detected were discarded due to potential debris contamination. These filtering steps resulted in 50,067 high quality nuclei in the final dataset. Finally, excitatory neurons, inhibitory neurons and non-neuronal cells were subset and reclustered using SCTransform workflow. Subclusters were further annotated using a human brain (middle temporal gyrus of the temporal cortex) snRNA-seq data [95] with high granularity. Overlap between the markers identified in the subclusters and reference datasets was performed by a Fisher’s exact test enrichment analysis as previously done [96, 97]. Clusters were annotated based on the highest enrichment score. We identified 12 cell-types in excitatory neurons, 17 in inhibitory neurons, and 12 in non-neuronal cells.

### Linking gene expression and gene regulatory elements with scMultiMap

To link gene expression with accessible chromatin, scMultiMap (v1.0.1) [27] was used. For each major cell class, we selected the top 5,000 most variable genes and the top 150,000 most variable peaks. Gene–peak pairs separated by ≤500kb were retained for downstream analysis. Associations were evaluated within each cell class separately by testing peak–gene pairs for significant correlation. In excitatory neurons: 340,129 gene–peak pairs were tested. After filtering (FDR<0.05), 20,901 pairs remained, 6,317 of which had non-zero correlations. In inhibitory neurons, 354,863 pairs tested; 78,820 survived FDR filtering; 11,343 had non-zero correlation. In non-neurons, 371,570 pairs tested; 101,130 passed FDR filtering; 34,145 showed non-zero correlation. In excitatory neurons, applying the above criteria yielded 2,869 significant pairs under control conditions and 3,448 under stimulation. In inhibitory neurons, 5,321 peak–gene links were found in controls versus 6,022 during stimulation. In glial cells, 17,475 links were detected under control conditions compared to 16,670 upon stimulation.

### Differential gene expression analysis

To identify differentially expressed genes (DEGs) between stimulation and control conditions in snRNA-seq data, we performed a pseudobulk analysis within each cell subtype. Briefly, raw counts were aggregated by sample using the R function AggregateExpression in Seurat. The resulting pseudobulk count matrices, one for each cell type, were analyzed using the DESeq2 package (v1.44.0) [98], which fits a Gamma-Poisson generalized linear model. We applied a gene expression filter, retaining only genes expressed in at least 10 samples. The DESeq2 model included relevant biological and technical covariates such as Age, Sex, Hemisphere, Epilepsy Duration, and number of nuclei. Genes were considered significantly differentially expressed if they had an adjusted p-value (padj) < 0.05 and an absolute log2 fold change > 0.3.

### Differential chromatin accessibility analysis

To identify differentially accessible regions (DARs) between stimulation and control conditions in snATAC-seq data, we conducted a pseudobulk analysis within each cell subtype. Specifically, raw chromatin accessibility counts were aggregated by sample using the AggregateExpression function in Seurat, resulting in pseudobulk count matrices for each cell type. These matrices were then analyzed using the DESeq2 package (v1.44.0), which fits a Gamma-Poisson generalized linear model. We applied a filter to retain only peaks accessible in at least 10 samples. The DESeq2 model incorporated relevant biological and technical covariates, including Age, Sex, Hemisphere, Epilepsy Duration, and number of nuclei. Peaks were considered significantly differentially accessible if they had an adjusted p-value (padj) < 0.05 and an absolute log2 fold change > 0.3.

### Gene ontology analyses

The functional annotation of the identified DEGs was performed using scToppR (v0.99.1) [99]. Enrichment was further confirmed using the R package clusterProfiler (v4.12.2) [100]. Significant categories were selected by a Benjamini-Hochberg corrected p-value < 0.05.

### Cell perturbation analyses

To identify which cell type was more perturbed by the stimulation, we used Augur (v1.0.3) [24] with linear regression test. The perturbations were further confirmed with MELD (v1.0.2) [101]. Standard parameters were used in both analyses.

### Cell-type specific gene regulatory networks with SCENIC+

We implemented the SCENIC+ (v1.0a1) [102] and pycisTopic workflows to construct gene regulatory networks (GRNs) for excitatory, inhibitory, and non-neuronal cell types. In this analysis, a GRN consists of multiple enhancer-driven regulons. Each regulon consists of one primary TF, its target chromatin regions (i.e., peaks containing the TF binding motif), and the target genes connected with those regions.

After generating a cistopic object using the preprocessed ATAC data, a Collapsed Gibbs Sampler was applied to model topics. For each subset, models with 10, 20, and 30 topics were trained and evaluated using pycisTopic’s built-in metrics. The model with 20 topics was selected as optimal. To identify candidate enhancer regions, we used three complementary approaches: 1) binarization of topics using the Otsu method, 2) selection of the top 3000 representative regions per topic to create region sets, and 3) identification of the top differentially accessible regions for each cell type, calculated by imputing region accessibility followed by a Wilcoxon test (log2 fold change > 0.3 and Benjamini–Hochberg-adjusted p-value < 0.05). These region sets were used as inputs for the SCENIC+ pipeline, which was run with standard parameters. Pycistarget and motif enrichment analysis using the Discrete Element Method (DEM) were then applied to determine whether the candidate enhancers were linked to specific transcription factors (TFs).

Next, regulons, defined as TF-region-gene triplets that include a specific TF, the set of regions enriched for its annotated motif, and the genes linked to those regions, were identified using a SCENIC+ wrapper function with default parameters. For each regulon, enrichment scores (AUC) of target genes in Stim and Control conditions were calculated using the AUCell algorithm via the score_eRegulons function. To identify cell-type-specific regulons, we computed RSS between cell types for Stim and Control using function regulon_specificity_scores. Regulons containing more than ten target genes were included in the analysis. We retained all regulons showing either positive or negative correlations between regions and genes, or between TF and genes. Additionally, we incorporated extended regulons, defined as those in which primary transcription factors are linked to target regions or genes based on orthologous annotations from non-human systems, when no direct annotations from human datasets were available. [29]

## Acknowledgments

S.S, B.G., and S.B. are supported by the CNDD Genomics and Bioinformatics Core at MUSC (NIH grant P20GM148302) and by the Biorepository & Tissue Analysis Shared Resource, Hollings Cancer Center, Medical University of South Carolina (P30 CA138313). This work was supported by NIH grants to H.M. (NS132443) and B.L. and G.K. (NS126143).

